# Stress-dependent activation of PQM-1 orchestrates a second-wave proteostasis response for organismal survival

**DOI:** 10.1101/2025.03.11.642454

**Authors:** Laura Jones, Valeria Uvarova, Daniel O’Brien, Holly McIntyre, Natalie R. Cohen, Robert H. Dowen, Patricija van Oosten-Hawle

**Author notes:** Corresponding author: Dr. Patricija van Oosten-Hawle. Contributed equally (Laura Jones, Valeria Uvarova).

## Abstract

Stress responses are controlled by specialized stress-responsive proteostasis transcription factors that rapidly upregulate protein quality components to re-establish protein homeostasis and safeguard survival. Here we show that the zinc finger transcription factor PQM-1 is crucial for stress survival in response to thermal and oxidative challenges. We provide mechanistic insight into the regulation of PQM-1 during stress that depends on ILS-DAF-16 signaling, as well as phosphorylation on threonine residue 268 that is located within a conserved AKT motif. Our data show that in reproductively mature adults and during well-fed conditions, PQM-1 induction requires DAF-16 and occurs during the recovery period post heat shock. Moreover, PQM-1 co-localizes with DAF-16 in the nucleus during the stress recovery phase. This regulatory dependency on DAF-16 is bypassed under dietary restriction, allowing PQM-1 to promote stress resilience independent of the ILS pathway. During both conditions, PQM-1 is crucial for the upregulation of cytosolic and endoplasmic reticulum stress response genes required for organismal recovery and stress resilience. Our transcriptional and bioinformatic analysis reveals that PQM-1 regulates a distinct set of target genes during the stress recovery phase, suggesting that PQM-1 may be involved in vital secondary wave stress response. Thus, our findings uncover a previously unrecognized mechanism of stress-dependent PQM-1 activation that integrates multiple environmental cues to ensure proteostasis and organismal survival.

## Introduction

All cells within an organism experience environmental or intrinsic proteotoxic stresses throughout life. To maintain and re-establish proteostasis, cells employ highly conserved stress response mechanisms that facilitate the upregulation of protein quality control components involved in protein folding and degradation ^1–3^, utilizing stress responsive transcription factors, including the heat shock factor 1, HSF-1, and fork head (FOXO) family members. In *C. elegans*, DAF-16 is the sole FOXO transcription factor and together with HSF-1, it is vital to promote resistance to a variety of stresses, such as heat ^4^, oxidative stress^5^ and pathogenic bacteria ^6^. Both HSF-1 and DAF-16 control the expression of molecular chaperones, including heat shock proteins, that are crucial for survival during environmental challenges as well as aging and are negatively regulated via the insulin-like signaling (ILS) pathway ^7,8^.

Recent evidence has shown that the zinc finger transcription factor PQM-1 takes an important role in the regulation of proteostasis maintenance during stress and aging, in addition to HSF-1 and DAF-16 ^9–12^. PQM-1 was originally discovered as a gene upregulated in response to oxidative stress in *C. elegans* ^13^ and identified to play a key role in the regulation of ILS-mediated longevity ^9^ and hypoxia^12^. As a lifespan regulating transcription factor, PQM-1 is thought to function in a mutually antagonistic manner with DAF-16 to control developmental gene expression by binding to genes harboring DAF-16 associated elements (DAE) ^9^. Moreover, PQM-1 is important for the developmental transition to reproductive adulthood together with the homeobox transcription factor CEH-60/PBX, by co-repressing DAE stress-response and longevity genes during adulthood in favor of gene networks required for reproduction, including fat metabolism and vitellogenesis related genes ^14,15^. However, how PQM-1 is regulated or activated following stress conditions has not been investigated in a comprehensive manner in adult *C. elegans*.

Here, we examined the PQM-1 response to a variety of environmental challenges during adulthood, including heat stress, oxidative stress and dietary restriction. Surprisingly, we find that PQM-1 and DAF-16 do not function in a mutually exclusive manner in response to heat stress, but that PQM-1 expression and nuclear accumulation depends on the presence of DAF-16. Importantly, in adults PQM-1 translocates to the nucleus only during the stress recovery phase, where it also co-localizes with DAF-16. Under dietary restriction, however, PQM-1 expression is uncoupled from DAF-16 regulation and insulin-like signaling to promote survival independently. Transcriptional analysis of *pqm-1* and *daf-16* loss-of-function mutants during a crucial timepoint in the stress recovery phase, combined with bioinformatic analysis of previous PQM-1 ChIP-seq datasets, revealed PQM-1 targets that are induced during stress recovery in both well-fed and dietary restricted animals and required for organismal stress resilience. Collectively, our findings suggest that PQM-1 might be involved in a “secondary wave” stress response that acts delayed but is complementary to the immediate transcriptional response orchestrated by DAF-16 and HSF-1.

## Results

### PQM-1 is a stress-responsive transcription factor, localizing to the nucleus after heat- and oxidative stress

We have previously shown that PQM-1 is required for heat stress survival in *C. elegans,* independent of the master regulator of the HSR, heat shock transcription factor (HSF-1) ^11^. PQM-1 is localized to intestinal nuclei in developing larvae during normal growth conditions, and becomes undetectable as soon as reproductive maturity in adulthood is reached ^9,11,14^. To analyze the relevance of PQM-1 activity during heat shock in larvae and adult animals we took advantage of a strain expressing a PQM-1::GFP::FLAG fusion protein. We monitored PQM-1::GFP::FLAG nuclear localization in Day 1 adults immediately after a 1-hour HS at 35C° (t = 0 hrs) and 2 hours post HS (+2 hrs) **(Fig. 1 A)**. PQM-1::GFP::FLAG is not visible in Day 1 adults during normal growth conditions **(Fig. 1A)**, or immediately after HS but becomes localized to the nucleus within 2 hours post HS **(Fig. 1A; Supp. Fig. 2A).** Nuclear localization reaches a peak between 1 and 3 hours post HS with 98% of PQM-1 being in the nucleus at the 2-hour post HS timepoint (P<0.0001) **(Fig. 1B, Supp. Fig. 2A)**. PQM-1 then exits the nucleus with only 12% of PQM-1 localized to the nucleus 6 hours post HS **(Fig. 1B, Supp. Fig. 2A)**. Previous research indicated that PQM-1 localizes to the cytosol during normal growth conditions (20°C)^9,11,14^. However we were unable to detect PQM-1::GFP fluorescence in Day 1 adults at basal conditions **(Fig. 1A; Supp. Fig. 2A)**, with both *pqm-1* mRNA **(Supp. Fig. 2C)** and protein expression (**Fig. 1C**) also being at low levels. Following HS, *pqm-1* mRNA transcripts were induced 5-fold **(Supp. Fig. 2C)**, with PQM-1 protein expression levels induced during the recovery period post HS, peaking between 1 and 3 hours (**Fig. 1C**). These findings indicate that, under physiological conditions, PQM-1 is expressed at very low levels in adult animals but becomes induced in response to stress. In contrast, PQM-1 is clearly visible in intestinal nuclei of L4 larvae, indicating transcriptional activity during development at normal growth conditions as shown before^9,11^. Similar to adults, PQM-1 localizes to the nucleus upon HS in L4 larvae within 2-hours post HS (**Suppl. Fig. 1A).**

**Figure 1.**
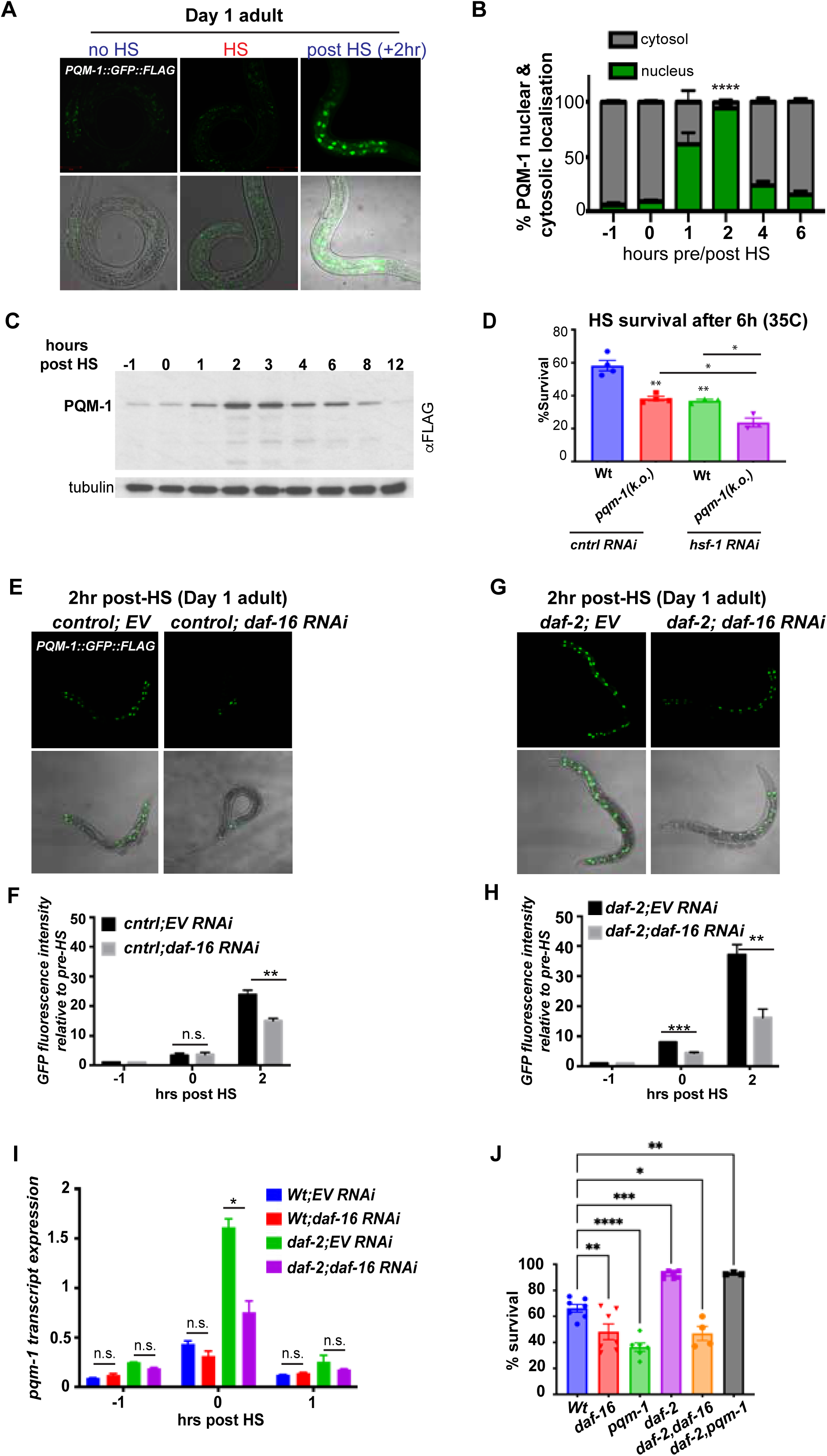
PQM-1 is a stress-responsive transcription factor in a ILS- and DAF-16 dependent manner. **(A)** Confocal images of PQM-1::GFP expression in Day 1 adult nematodes grown at 20°C before HS (− 1hr), immediately after a 1-hour 35°C HS (0 hr) and 2 hours post HS (+ 2 hr). PQM-1::GFP is not visible in intestinal nuclei in adults during normal growth conditions or HS, but appears in the nucleus during recovery. Scale bar = 50 μm. **(B)** Quantification of PQM-1 nuclear and cytosolic localization before (−1hr), immediately after HS (0 hr) and recovery post HS (1 hr – 6 hr post HS). n > 20 per condition; *P<0.05. Error bars represent ± SEM. **(C)** Western blot analysis of PQM-1::GFP::FLAG expression before (−1hr), immediately after HS (0 hr) and recovery post HS (1 – 12 hrs), using an anti-FLAG antibody. Tubulin was used as a loading control. **(D)** Heat stress survival of Day 1 N2 Bristol (Wt) and *pqm-1(ko)* mutant nematodes during control RNAi or *hsf-1* RNAi after a 6-hour HS at 35°C. Survival was scored after a 16 hr recovery period at 20°C. n = 20; three biological replicates. Bar graph represent SEM, **P < 0.01; *P<0.05 **(E)** PQM-1::GFP localization to intestinal nuclei in Day 1 adult control animals (strain OP201) after a 2 hour recovery post HS; and **(F)** PQM-1::GFP nuclear localization in *daf-2* mutants after a 2-hr recovery post HS, either grown on control RNAi (EV) or *daf-16* RNAi. **(F &H)** Quantification of GFP fluorescence intensity of PQM-1 localized to intestinal nuclei immediately after HS (0 hr) and 2 hours post HS, relative to fluorescence intensity levels before HS (−1hr) in **(F)** control animals and **(H)** *daf-2* mutants. Nematodes were grown on control (EV) or *daf-16* RNAi before HS treatment. **(I)** *pqm-1* transcript expression levels of control (wt), *daf-2* mutants grown on control (EV) or *daf-16* RNAi before (−1hr), after HS (0hr) and 1-hour post HS (+1hr). **(F, H, I)**. Student’s t-test was performed to calculate statistical significance of GFP fluorescence intensity or transcript expression after HS relative to levels before HS (20°C). *P < 0.05; **P < 0.01; ***P < 0.001; n.s. = not significant. **(J)** Heat stress survival rates of age-synchronized Day 1 adult *daf-16(mu86)* and *pqm-1(ko)* mutants in a control- or *daf-2* mutant background, after a 6-hour HS at 35°C. Bargraph represents S.E.M. *P < 0.05; ***P < 0.001, ****P < 0.0001.

We next investigated the PQM-1 response in adults exposed to oxidative stress via treatment with 100 mM paraquat for 1 hour **(Supp. Fig. 1B)**. Similar to heat stress, PQM-1::GFP::FLAG protein expression levels **(Supp. Fig. 1C)**, as well as *pqm-1* transcript levels (**Supp. Fig. 1D)** were induced and localized to the nucleus following exposure to oxidative stress. Together, these observations demonstrate that PQM-1 is a stress-inducible transcription factor in adults that accumulates in the nucleus only during the recovery phase post stress, whereas in developing larvae it is constitutively induced and present in the nucleus at physiological conditions.

Given that PQM-1 expression and nuclear localization in adult *C. elegans* appears to be stress-dependent, we investigated whether PQM-1 is essential for *C. elegans* survival to heat and oxidative stress conditions. We found that *pqm-*1 null mutants (*pqm-1(k.o.*)) exhibited reduced heat stress survival rates, similar to nematodes treated with *hsf-1* RNAi (**Fig. 1D**; 40%, P < 0.01)^11^. Notably, the low survival rate of *pqm-1*(k.o.) mutants further decreased to 25% (P<0.05) upon *hsf-1* knock down by RNAi (**Fig. 1D**), indicating that heat stress survival is mediated by PQM-1 in an *hsf-1* independent pathway, as previously observed ^11^. Under oxidative stress *pqm-1(k.o.)* mutants also displayed reduced survival (60%, P<0.05) comparable to *C. elegans* treated with *skn-1* RNAi, one of the major transcription factors regulating the response to oxidative stress (Supp. Fig. 1E) ^16,17^. Depletion of both *skn-1* and *pqm-1* further lowered survival, albeit not statistically significant (50%, P=0.07; **Supp. Fig. 1E**). In summary, these findings demonstrate that PQM-1 is a stress-induced protein that is crucial for survival under both heat and oxidative stress conditions.

### DAF-16 regulates PQM-1 expression after heat stress

We next examined how PQM-1 is regulated during heat shock (HS) in adults. Nuclear localization of DAF-16 and PQM-1 is thought to be mutually exclusive, governed by the insulin-like signaling (ILS) pathway and HS^9^. Because nuclear DAF-16 is known to restrict the nuclear localization of PQM-1, we hypothesized that reducing *daf-16* expression would enhance PQM-1 nuclear localization during HS^9^. Surprisingly, however, *daf-16* RNAi decreased PQM-1 nuclear localization following HS (**Fig. 1E**), resulting in a 30% reduction in nuclear PQM-1::GFP fluorescence intensity (**Fig. 1F**). In heat-shocked *daf-2* mutants, which exhibit elevated DAF-16 activity, *pqm-1* transcript levels (**Fig. 1I**) and nuclear PQM-1::GFP intensity (**Fig. 1G, 1H**) both increased two-fold relative to wild type. Conversely, RNAi-mediated knockdown of *daf-16* in *daf-2* mutants lowered PQM-1::GFP intensity by 50% (**Fig. 1G, 1H;** P < 0.05) and reduced *pqm-1* transcript expression after HS also by 50% (**Fig. 1I**, P < 0.05). Consistent with these transcript levels, PQM-1 protein expression also declined upon *daf-16* RNAi in control and *daf-2* mutants (**Fig. 2E and 3F**). These findings indicate that PQM-1 transcriptional regulation during HS depends on DAF-16 and that both transcription factors may be required for robust heat stress survival. Consistent with these findings, *C. elegans* lacking either *daf-16* or *pqm-1* exhibit ∼60% lower survival under heat stress (**Fig. 1J**), paralleling the diminished induction and nuclear localization of PQM-1 during *daf-16* RNAi. By contrast, the enhanced heat stress resistance of *daf-2* mutants appears to be independent of *pqm-1* and relies solely on *daf-16* (**Fig. 1J**). Overall, the results show that DAF-16 promotes the accumulation of PQM-1 protein during HS, while the increased survival of *daf-2* mutants is not contingent on PQM-1.

**Figure 2.**
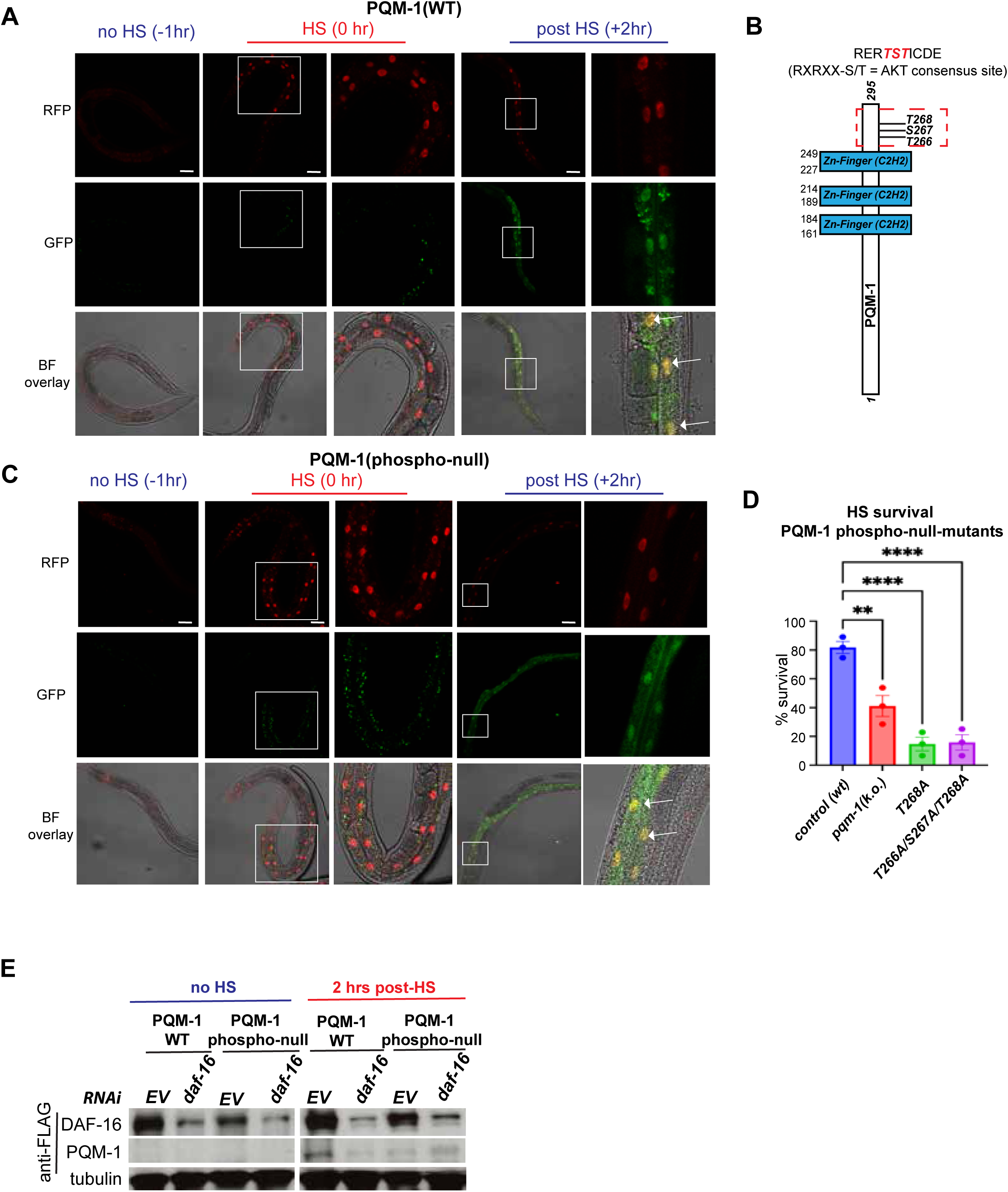
PQM-1 and DAF-16 co-localize in the nucleus after HS. Confocal images of nuclear localization of DAF-16 (mKate2) and PQM-1 (GFP) in **(A)** wild type PQM-1 (Biotag::GFP::3xFLAG::PQM-1; strain DLS608) before (−1hr), immediately after HS (35°C; 0hr) or 2 hours post HS. White arrows indicate co-localization of GFP::PQM-1 and DAF-16::mKate2 at the +2-hr timepoint. Scale bar = 25 μm. (**B)** Schematic of the PQM-1 domain structure, highlighting the three zinc finger domains and the C-terminal phosphorylatable RERTST motif. **(C)** Confocal images of nuclear localization of DAF-16 (mKate2) and PQM-1 (GFP) in the *pqm-1(phospho-null)* mutant (Biotag::GFP::FLAG::PQM-1(T266A/S267A/T268A); strain DLS610) before (−1hr), immediately after HS (35°C; 0hr) or 2 hours post HS. White arrows indicate co-localization of GFP::PQM-1 and DAF-16::mKate2 at the +2-hr timepoint. Scale bar = 25 μm. **(D)** Heat stress survival rates of age-synchronized Day 1 adult pqm-1(ko) mutant and PQM-1 phospho-null mutants T268A and T266A/S267A/T268A, after a 6-hour HS at 35°C. n=3; Bargraphs represent S.E.M. *P < 0.05; ****P < 0.0001. **(E**) Western blot analysis PQM-1 and DAF-16 expression at 20°C and 2 hrs post HS in Day adult wildtype PQM-1 (strain DLS608) and *pqm-1(phospho-null)* mutant (strain DLS610) during control (EV) and *daf-16* RNAi.

**Figure 3.**
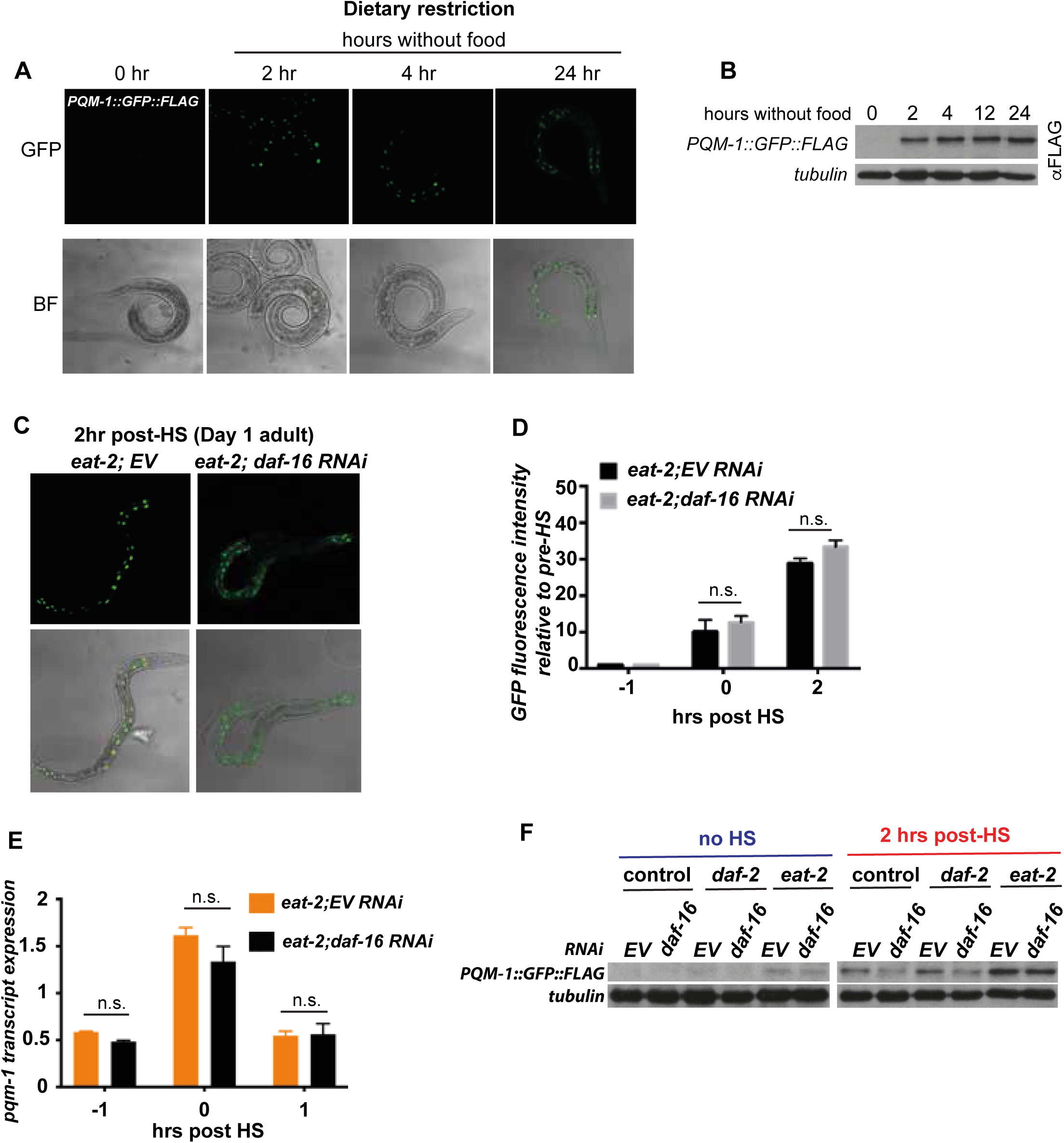
DR promotes stress resistance via PQM-1, independent of DAF-16. **A)** PQM-1::GFP::FLAG in Day 1 adults is localized to intestinal nuclei after a 2-, 4-, or 24 hr period of starvation. PQM-1 is not detectable in nuclei of well-fed Day 1 adult nematodes (0 hr). **(B)** Western blot analysis and quantification of PQM-1::GFP::FLAG expression levels in Day 1 adults before (0 hrs) and during 2, 4, 12 and 24 hours of starvation, using an anti-FLAG antibody. Tubulin was used as a loading control. **(C)** PQM-1::GFP::FLAG nuclear localization in *eat-2* mutants after a 2-hr recovery post HS, and grown on control (EV) or *daf-16* RNAi. **(D)** Quantification of GFP fluorescence intensity of PQM-1 localized to intestinal nuclei immediately after HS (0 hr) and 2 hours post HS, relative to fluorescence intensity levels before HS (−1hr) in *eat-2* mutants either grown on control (EV) or *daf-16* RNAi. **(E)** *pqm-1* transcript expression levels of *eat-2* mutants grown on control (EV) or *daf-16* RNAi before (−1hr), immediately after HS (0hr) and 1-hour post HS (+1hr). **(F)** Western blot analysis of PQM-1::GFP::FLAG expression levels in control (strain OP201), *daf-2* and *eat-2* mutants before (no HS) and 2 hours post HS and treated with control (EV) and *daf-16* RNAi..

### PQM-1 and DAF-16 co-localize in the nucleus post HS, independent of PQM-1 phosphorylation

Because PQM-1 expression appears to be dependent on DAF-16 during HS, we examined whether this is reflected in their nuclear localization dynamics using a CRISPR/Cas9-edited strain expressing endogenously tagged versions of mKate2::3xFLAG::DAF-16^18^ and Biotag::GFP::3xFLAG::PQM-1. Immediately following a 1-hour HS at 35°C (time = 0 hrs), DAF-16 is localized to the nucleus, whereas GFP::PQM-1 is not detectable (**Fig. 2A)**. Interestingly, 2-hours post HS both PQM-1 and DAF-16 co-localize simultaneously in the nucleus (**Fig. 2A**) suggesting that DAF-16 must enter the nucleus first, followed by the production and nuclear localization of PQM-1. This indicates that PQM-1 might be involved in a delayed transcriptional program initiated in the recovery phase after stress, whereas DAF-16 is involved in an immediate response.

To further investigate how PQM-1 is regulated we focused on post-translational modifications. PQM-1 contains a RERTST motif at the C-terminal end, representing a consensus AKT or AGC kinase phosphorylation site, that was previously proposed to be important for its DNA binding activity (**Fig. 2B**)^14,15^. We therefore questioned whether PQM-1 phosphorylation at this conserved motif was important for heat stress survival. We examined two PQM-1 phospho-mutants: T268A and a triple phospho-(null)-mutant T266A/S267A/T268A where all possible phosphorylation sites of the RERTST motif were mutated. Surprisingly, the heat stress survival rate of the PQM-1 T268A mutant (15%, P < 0.0001) and the triple mutant T266A, T267A, T268A (15.8 %; P<0.0001) were even lower than the *pqm-1(k.o.)* deletion mutant compared to the control (40%; P < 0.01) (**Fig. 2D**). These data indicate that phosphorylation at T268 is essential for HS survival and that mutating these sites produces an anti-morphic phenotype, emphasizing the importance of the RERTST motif for the stress-protective function.

We next asked whether the reduced survival rate of the *pqm-1* phospho-mutants stemmed from impaired nuclear localization or accumulation of the transcription factor after HS. Unexpectedly, the heat-stress sensitive PQM-1 phospho-null-mutant T266A/S267A/T268A is not impaired in its ability to localize to the nucleus **(Fig. 2C)**. Both PQM-1(WT) and the (TST)phospho-mutant show clear co-localization with DAF-16 in the nucleus (**Fig. 2A and Fig. 2C**). Taken together, these results demonstrate that DAF-16 nuclear localization precedes PQM-1 expression, supporting the idea that DAF-16 is required to induce PQM-1. Furthermore, DAF-16 and PQM-1 do not act in a mutually exclusive manner but instead co-localize in the nucleus during HS recovery, regardless of the PQM-1 phosphorylation status at the RERTST motif.

Because our data suggests that *daf-16* is required for the induction of PQM-1 protein during HS recovery, we tested this further by *daf-16* RNAi in the GFP::PQM-1 wild type and phospho-null-mutant. Depletion of *daf-16* by RNAi reduced PQM-1(WT) and PQM-1(TST-phospho-null) protein levels expression levels after HS (**Fig. 2E**), further supporting the observation that PQM-1 expression depends on DAF-16.

### Dietary restriction uncouples PQM-1 from DAF-16 control

PQM-1 activity was previously shown to be regulated by dietary restriction (DR)^10^. DR is one of the most powerful interventions to increase stress resistance and can be mediated in an ILS/DAF-16 activity dependent manner ^19^ as well as independent manner ^20–23^. We therefore asked whether DR regulates PQM-1 expression and activity dependent or independent of the ILS pathway to promote *C. elegans* stress resistance. First, we examined two DR regimens: a short DR initiated for 24 hours in well-fed Day 1 adults and a chronic DR regimen throughout life using the *eat-2*(*ad453*) mutation, which reduces pharyngeal pumping rates and consequently food intake^24^. After a 24-hour bacterial deprivation regimen that was started in Day 1 adults, PQM-1 nuclear localization (**Fig. 3A**) as well as protein expression are increased and the higher PQM-1 expression levels are sustained during 2, 4, 12 and 24 hours of starvation **(Fig. 3B)**.

We then questioned whether PQM-1 expression and nuclear localization is influenced by DAF-16 during reduced food intake using the *eat-2(ad453)* mutant. Surprisingly, *daf-16* RNAi in *eat-2* mutants exposed to heat shock did not significantly reduce nuclear localization of PQM-1 **(Fig. 3C)** or nuclear PQM-1::GFP fluorescence intensity **(Fig. 3D)**. Consistently, *pqm-1* transcripts (**Fig. 3E)** and PQM-1 protein levels (**Fig. 3F**) remained unchanged by *daf-16* RNAi in the *eat-2* mutant. The induced PQM-1 protein expression levels in *eat-2* mutants, even in the absence of HS, remained unaltered upon *daf-16* RNAi **(Fig. 3F)**. In contrast, *daf-16* RNAi in *daf-2* mutants decreased PQM-1 expression (**Fig. 3F**). This indicated that PQM-1 expression can be uncoupled from DAF-16 during DR. This observation is further corroborated by the 50% reduced heat stress survival of *daf-2* mutants in a *daf-16* loss of function mutant (*daf-16(mu86)*), whereas *daf-2;pqm-1* double mutants remained unaffected (∼100% survival rate) **(Fig. 1J)**. In contrast, during DR, *eat-2;daf-16* double mutants are unaffected by the loss of *daf-16*, whereas the increased survival rates of the *eat-2* mutant is reduced by 50% upon *pqm-1* depletion **(Fig. 4I)**. Both, the *eat-2;pqm-1* double mutant **(Fig. 4I)** and the *pqm-1* null mutant alone **(Fig. 1J)** showed similar survival rates following HS (∼50%, P < 0.01), indicating that DR-induced signaling regulates PQM-1 activity independently from the ILS/DAF-16 signaling pathway. In summary, this data demonstrates that during well-fed conditions PQM-1 expression depends on ILS and DAF-16. During dietary restriction, as in the *eat-2* mutant, PQM-1 expression is uncoupled from ILS/DAF-16 signaling, depending on a, yet to be identified, different upstream signaling cue that regulates its expression.

**Figure 4.**
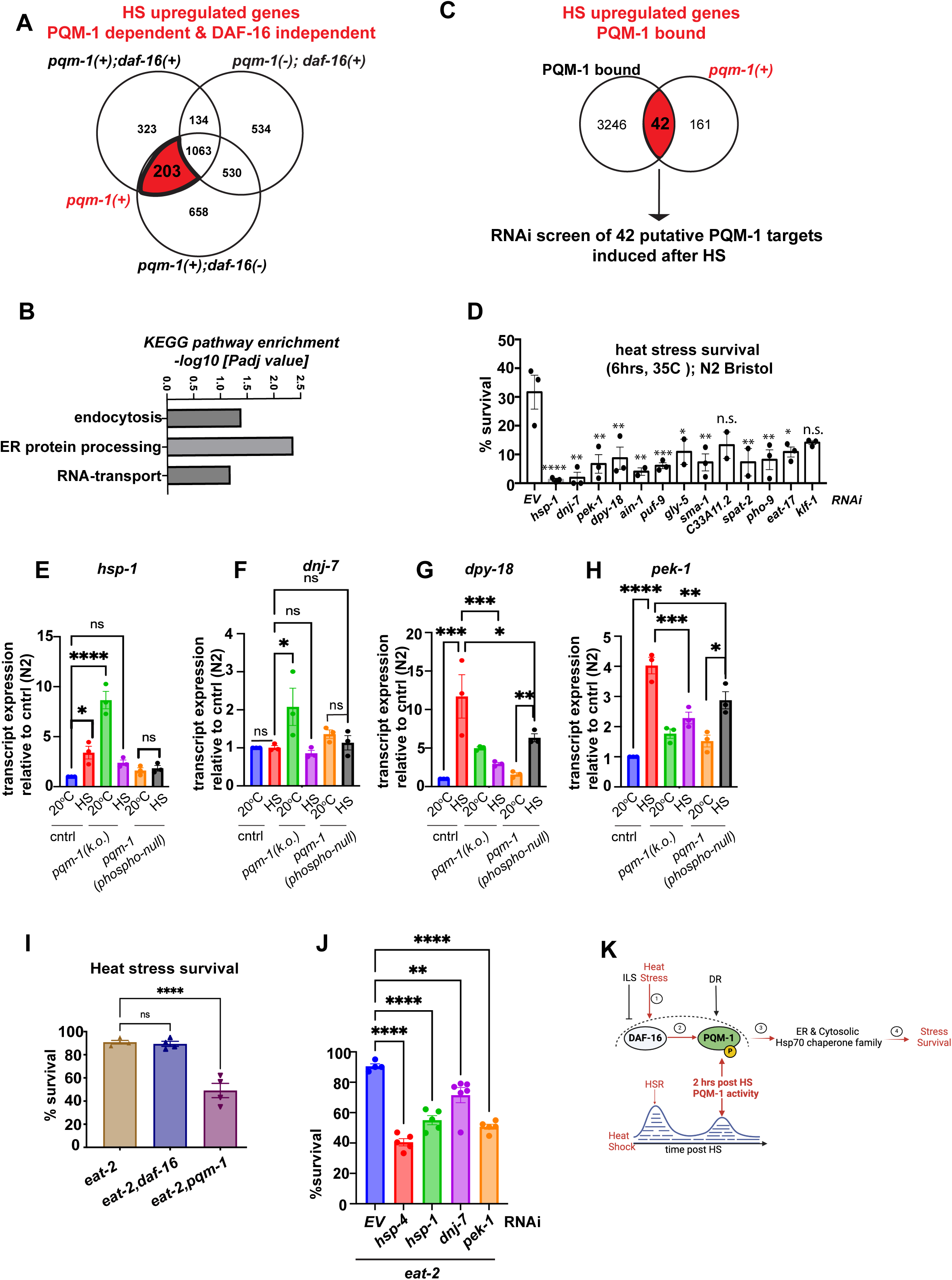
PQM-1 regulates ER resident and cytosolic chaperones in response to heat stress. **(A)** Venn-diagram analysis of upregulated genes 2 hours post HS to identify genes dependent on the presence of PQM-1, but independent of DAF-16. The red-shaded area represents 203 genes dependent on *pqm-1*. **(B)** KEGG pathway enrichment categories of the 203 genes are shown as the −log10 transformation of the Benjamini-Hochberg corrected p-value. **(C)** Venn-diagram analysis of HS-upregulated genes that are directly bound by PQM-1 in L3 larvae according to previous PQM-1 ChIP-Seq analysis (Niu et al., 2011; Dowen 2019). **(D)** Heat stress survival rates of wild-type (N2) animals after RNAi-mediated knockdown of 13 PQM-1-dependent HS-induced genes. Statistical analysis one-way ANOVA. n=3; Related to Suppl. Figure 3F. **(E-H)** Transcript expression of *pqm-1*-dependent chaperone and stress genes *hsp-1* **(E)**, *dnj-7* **(F)**, *dpy-18* **(G)**, *pek-1* **(H)**, in control (N2), *pqm-1(ko)* mutant and *pqm-1 (phospho-null)* mutant before and after HS (1h, 35°C). n=3. Error bars represent S. E. M. Statistical analysis one-way ANOVA. *P<0.05; **P<0.01; ***P<0.001; ****P<0.0001. **(I)** Heat stress survival rates of age-synchronized Day 1 adult *daf-16(mu86)* and *pqm-1(ko)* mutants in a control, or *eat-2(ad453)* mutant background, after a 6-hour HS at 35°C. **(J)** Heat stress survival rates of age-synchronized Day 1 adult eat-2 mutants after RNAi-mediated knockdown of *hsp-4, hsp-1, dnj-7*, and *pek-1.* n≥3; one-way ANOVA. **P<0.01, ****P < 0.0001; n.s. = not significant. Error bars represent S.E.M. **(K)** Working model of PQM-1 regulation after heat shock in adults. Following heat stress, DAF-16 localizes to the nucleus (1) to regulate the expression of immediate stress response genes. (2) During stress recovery, 2-hours after HS, PQM-1 then accumulates into the nucleus and co-localizes with DAF-16. During dietary restriction (DR), regulation of PQM-1 expression becomes independent of DAF-16 to regulate stress survival. PQM-1 transcriptional activity depends on phosphorylation on its RERTST motif, which is (3) required for transcription of ER resident Hsp70 family chaperones (*dnj-7, dpy-18, hsp-4*) and cytosolic Hsp70 (*hsp-1*), to promote stress survival (4).

### PQM-1 promotes ER proteostasis genes required for DR-induced stress survival

To elucidate the molecular basis underlying PQM-1 function in stress survival, we next analyzed transcriptional gene expression changes that occur during 2-hour timepoint post HS, when the nuclear localization of PQM-1 is at its highest. Specifically, we performed mRNA -Seq of *pqm-1(ko)* mutants 2 hours post HS compared their expression profiles with those of wild type animals (N2) and the *daf-16(mu86)* knockout mutant (**Supp. Fig.3 A-C**). In *pqm-1(ko)* mutants, 2261 genes were upregulated in response to HS (**Supp. Fig. 3B**), compared to 1723 genes in wild type animals (**Supp. Fig. 3A**) and 2454 genes upregulated in *daf-16* mutants (**Supp. Fig. 3C**). Using Venn diagram analysis, we identified 203 genes (P <0.05) that are upregulated exclusively by PQM-1 after HS in a *daf-16*-independent manner (**Fig. 4A**; *red-shaded overlap area*). The KEGG pathway terms most strongly enriched among the 203 upregulated genes corresponded to endocytosis, ER protein processing and RNA transport (**Fig. 4B**). We next analyzed which of these 203 genes are directly bound by PQM-1 by comparing them to published PQM-1 ChIP Seq data sets, corresponding to the PQM-1 binding profile collected from L3 larvae^25^ and Day 1 adults^15^ during normal growth conditions (20°C). We identified 42 candidate genes that may be directly regulated by PQM-1 following HS. Among these genes were chaperones including heat shock cognate protein *hsp-1*/Hsc70, and two ER-localized chaperones, the DnaJ protein *dnj-7/*DnaJC3 and the TPR domain containing prolyl-4-hydroxylase *dpy-18*, and the ER stress sensor protein *pek-1*/PERK. We also examined chaperone gene induction in the wild type and *daf-16* mutant to evaluate their expression profile during stress recovery. Of the 170 known chaperone genes in *C. elegans* ^26^, 33 were induced in wild-type animals at the critical timepoint two hours after heat shock. Among these, which included *hsp-70, F44E5.4,* and *hsp-16.2* - were induced in wild type animals, neither depending exclusively on *pqm-1* or *daf-16*. This suggests dependence of these genes on the master regulator of the cytosolic heat shock response, HSF-1 (**Supp. Fig. 3D**). Only three chaperones *dnj-7, hsp-1* and *dpy-18*, as well as the ER stress sensor *pek-1* were upregulated in a *pqm-1*-dependent manner (**Supp. Fig. 3E**). Five chaperone genes were exclusively dependent on *daf-16* at the 2-hour post HS time point, that included two cytosolic Hsp70 isoforms (F44E5.4 and F44E5.5), ER resident chaperones of the Hsp70 chaperone- and co-chaperone family (T14G8.3 and Y73F8A.26, respectively) and a small heat shock protein (F08H9.4) (**Supp. Fig. 3D and 3E**). PQM-1 target genes recently identified to be involved in the hypoxia stress response, in particular *dod-17, dod-19* and *dod-24* ^27^ are downregulated following heat stress in our dataset, suggesting they could be repressed by PQM-1 during HS recovery.

To determine which of the 42 HS-induced putative PQM-1 target genes are required for heat stress survival, we performed a mini-RNAi screen utilizing wild type (N2) *C. elegans* and measured resistance to a 6-hour HS (35°C) (**Supp. Fig. 3F**). Genes whose knockdown lowered survival below 15% were classified as “hits,” revealing 13 candidates, that were re-examined for their requirement for heat stress survival (**Fig. 4D**). We then examined the induction of these 13 genes in *pqm-1(ko)* and *pqm-1(phospho-null)* mutants after a 1 hour HS at 35°C, compared with wild-type animals (**Supp. Fig. 3G**). Our results showed that *hsp-1, pek-1, dpy-18, pho-9* (a phosphatase), and *klf-1* (a zinc finger protein) are normally upregulated by HS, but their induction was reduced by at least 50% in both *pqm-1(ko)* and *pqm-1(phospho-null)* strains (**Supp. Fig. 3G**). These findings support earlier suggestions that *pqm-1* regulates a distinct network of chaperone genes^10^, although the mechanisms by which it confers stress resilience remained unclear. Interestingly, *hsp-1, dpy-18* and *pek-1* are induced in the control animals (N2) after HS (**Fig. 4E, 4G and 4H, respectively**), but not *dnj-7*, which is only induced in the *pqm-1(ko)* mutant at 20°C (**Fig. 4F**). *Hsp-1* transcripts are also induced 8-fold at the permissive temperature in the *pqm-1(ko)* mutant (**Fig. 4E**). This implies that *pqm-1* might act as a transcriptional repressor for both *hsp-1* and *dnj-7* under normal conditions, but is required for *hsp-1* induction after HS. In contrast, the *pqm-1(phospho-null)* mutant shows no induction of *hsp-1* and *dnj-7* after HS (**Fig. 4E and 4F**). Moreover, heat shock induction of *dpy-18, pek-1, pho-9* and *klf-1* was significantly reduced in the *pqm-1(phospho-null)* mutant compared to the N2 control (**Fig. 4G-4H, Supp. Fig. 3G**), indicating that phosphorylation of PQM-1 is necessary for their transcriptional activation. Thus, while phosphorylation does not seem to affect nuclear localization of PQM-1, this data suggests that it might be required for full transcriptional activity of PQM-1. Furthermore, the ER resident chaperones *dnj-7* and *dpy-18* as well as the ER stress sensor *pek-1* depend on *pqm-1*, underscoring a broader role for PQM-1 in being involved in both cytosolic (*hsp-1*) and ER related stress responses (*dnj-7, pek-1, dpy-18*). Interestingly, DR has previously been linked to increased proteostasis through ER hormesis ^28^. We therefore asked whether the increased stress resilience of *eat-2* mutants is mediated by *pqm-1*-dependent ER gene targets. Indeed, heat stress survival of *eat-2* mutants following RNAi-mediated knock down of the putative PQM-1 targets *dnj-7, pek-1, dpy-18* and the ER resident Hsp70 *hsp-4*/BiP was reduced by ∼50% **(Fig. 4J),** similar to *pqm-1* depletion in *eat-2* mutants **(Fig. 4I)**. This indicates that PQM-1 contributes to the increased stress resilience of *eat-2* mutants by regulating expression of these genes, albeit it may not be the sole transcription factor involved. Notably, after HS, regardless of upstream regulation by ILS/DAF-16 or DR, PQM-1 appears to govern a similar set of stress response genes that are induced at the 2-hour timepoint following HS to promote stress resilience (**Fig. 4D and 4J**). Together, these results suggest that *pqm-1* is induced and activated during stress recovery, and drives a distinct transcriptional program that may act as a “second wave stress response” crucial for organismal survival.

## Discussion

The adaptation to environmental challenges is mediated via stress-responsive transcription factors that regulate the expression of proteostasis effectors to promote survival during acute and chronic proteotoxic stress. Here, we provide new mechanistic insight into the regulation of the stress-responsive transcription factor PQM-1^9,11,12,27^ following a variety of environmental challenges including heat, oxidative stress and DR. Our data shows that in *C. elegans* adults PQM-1 expression is not induced immediately following stress, but within a two-hour time frame during stress recovery. This can be regulated through ILS/DAF-16-dependent mechanisms, in which both transcription factors co-localize in the nucleus during stress recovery, or independently of DAF-16 under DR conditions. PQM-1 transcriptional activity during this stress recovery period depends on phosphorylation on the conserved T268 site. We propose that the delayed peak of PQM-1 nuclear localization at two hours post HS represents a delayed “second wave” stress response when PQM-1 initiates a distinct transcriptional program involving ER and cytosolic resident chaperones crucial for stress resilience and survival (see **Fig. 4K** for a working model). This second wave appears to be separate from the immediate transcriptional stress responses mediated by DAF-16 and HSF1, and highlight a previously underappreciated layer of the heat shock response network involving coordination between multiple transcription factors.

One of the surprises of this study is that PQM-1 and DAF-16 both colocalize in the nucleus following heat stress, with DAF-16 nuclear localization preceding that of PQM-1. It was previously shown that PQM-1 and DAF-16 function antagonistically, with PQM-1 and DAF-16 localizing to intestinal nuclei in a mutual exclusive manner^9^. During developmental larval stages PQM-1 is in the nucleus and DAF-16 is localized primarily to the cytoplasm ^9^. Upon heat treatment, DAF-16 enters the nucleus, while PQM-1 leaves it, demonstrating opposite subcellular localizations as a function of time^9^. An important difference is that these previous experiments were conducted during larval stages and not during Day 1 of adulthood as in the present study. While PQM-1 is primarily localized to intestinal nuclei in developing larvae, its localization becomes more diffuse as *C. elegans* reaches reproductive maturity during young adulthood^11^. Environmental challenges such as heat, oxidative stress and dietary restriction during the adult stage induces PQM-1 protein expression and leads to its accumulation in the nucleus. Thus, under stress conditions during adulthood both PQM-1 and DAF-16 can coexist in the nucleus and may even jointly associate on certain gene regions. Indeed during cold stress, both PQM-1 and DAF-16 were shown to bind to the DAF-16 associated element (DAE) on the promoter of hypothermia-induced genes^29^, suggesting that one transcription factor recruits the other one to the promoter. Based on our observation that DAF-16 localizes to the nucleus before PQM-1 during heat stress, suggests that DAF-16 may not only control its expression but also associate with PQM-1 during this condition to promote survival. This can be a likely scenario, as PQM-1 was shown to associate with the CEH-60 and UNC-62 transcription factors at DAF-16 associated elements (DAE) to repress gene expression at the L3 larval stage^15^. This demonstrates that PQM-1 activation is differentially regulated by potential interaction with other transcription factors, dependent on developmental and environmental inputs. Such a differential regulation may become important as *C. elegans* transition from developing larva to reproductive adults, which is marked by a redistribution of intestinal lipids to the germline and oocytes. Indeed, PQM-1 is significantly involved in this transition step by activating fat metabolism genes required for reproduction together with the transcription factors CEH-60 and UNC-62^15^. Concurrently, in *C. elegans* adults PQM-1 is crucial to safeguard the expression and allocation of lipids to developing embryos under oxygen-limiting conditions^12^.

Our data indicates that PQM-1 activity is tightly regulated by multiple signaling pathways that respond to a diverse range of systemic and cellular stress, metabolic and developmental signals and suggests that fine-tuning of its transcriptional activity is critical to ensure organismal survival and stress resilience. Indeed, phosphorylation likely controls key aspects of PQM-1 transcriptional activity^15^. The PQM-1 TST-phospho-null mutant still localizes to the nucleus after heat stress, but shows reduced heat stress survival rates and reduced induction of PQM-1 target genes, suggesting de-phosphorylation at the RERTST motif sites impacts PQM-1 transcriptional activity but not nuclear localization. Further understanding of PQM-1 post translational modifications and how they impact protein activity will require more detailed biochemical exploration such as co-immunoprecipitation experiments in the future. Our finding that the *pqm-1(phospho-null)* mutation is more deleterious than complete absence of *pqm-1* itself is counterintuitive but noteworthy. It indicates that the substitution to either T268A (or the triple *phospho-null mutant*) could lead to an altered conformation of PQM-1 which not only disrupts its transcriptional activity, but also its interaction with other transcription factors involved in the same stress response. Thus, such a conformational change might impact transcriptional activity which converts the transcription factor from an active into a repressive state. This is at least in part supported by our data showing that induction of stress-related target genes in the *pqm-1(phospho-null)* mutant is significantly impaired, but not completely absent as for *dpy-18* and *pek-1* (Fig. 4E-4H). Such a mechanism could also account for the observed nuclear presence of the PQM-1 phospho-null mutant, despite a decrease in survival rates.

It is noteworthy that the basal expression and nuclear accumulation of PQM-1 is induced at the permissive temperature during DR conditions independently of DAF-16. Thus transcriptional control via DAF-16 is uncoupled during DR, which also promotes lifespan and proteostasis in a PQM-1 dependent manner ^9,10,14^. In addition to DR, signals from the germline also influence PQM-1 and DAF-16 activity, via neuronal gonadotropin signaling^10,30^. While our results show PQM-1 localization is regulated by dietary restriction, we cannot exclude that signals from the reproductive system also influence DAF-16 and/or PQM-1 localization and activation in response to stress to enhance proteostasis and longevity^10^. Another interesting observation is that while PQM-1 expression is elevated in a *daf-2* mutant following HS, it is at the same time dispensable for the increased thermotolerance of *daf-2* mutants. This indicates that *daf-16*, when active in the *daf-2* mutant, further promotes *pqm-1* expression after HS. Because *daf-16* itself is crucial for heat stress survival, it can at least in part compensate for the absence of *pqm-1* in the *daf-2* mutant. Moreover, the DAF-2/ILS pathway also regulates HSF-1 activity ^8,31^, which contributes to the sustained survival of *daf-2* mutants even in the absence of *pqm-1*.

PQM-1 activity is crucial for both heat- and oxidative stress survival in addition to HSF-1, DAF-16 and/or SKN-1. The nuclear accumulation of both DAF-16 and HSF-1 ^32,33^ occurs rapidly and precedes that of PQM-1 following stress. Indeed, *pqm-1* depletion in addition to *hsf-1* or *skn-1* knockdown further reduces stress survival, suggesting that *pqm-1* acts complementary to the *hsf-1* or *skn-1* mediated stress response. Thus, we propose that PQM-1 induces a stress-responsive gene network that functions as a “second wave” response that is complementary to the HSF-1, SKN-1 and DAF-16 promoted acute-stress transcriptional networks. This puts an additional layer on the dynamics of transcriptional responses to stress that considers multiple transcription factors involved in the regulation of proteostasis and aging and that warrants further exploration in the future.

## Material and Methods

### Nematode strains and growth conditions

*C. elegans* were grown on NGM Agar plates seeded with *E.coli* strain OP50-1 and cultured using standard methods, as described in (Brenner, 1974). Strains were obtained from the Caenorhabditis Genetic Center (CGC). For a detailed description of *C. elegans* strains generated in this study, please refer to Supplemental Table 1.

### RNAi experiments

For RNAi-mediated knockdown of indicated genes, adult nematodes were allowed to lay eggs overnight on 60 mm NGM plates seeded with *E. coli* strain HT115(DE3), transformed with the appropriate RNAi vector (J. Ahringer, University of Cambridge, Cambridge, UK).

### Confocal microscopy and quantification of fluorescence intensity

Day 1 adult *C. elegans* nematodes were immobilized on 1% agarose pads in 5 mM levamisole. Nematodes were imaged using a Zeiss LSM880 confocal microscope and X 20 objective. GFP expression was quantified in the 4-5 posterior intestinal cells using ImageJ.

### Quantification of PQM-1::GFP nuclear localization

*C. elegans* were age-synchronized by egg laying and >25 Day 1 adults were exposed to HS by incubating in a water bath equilibrated at 35°C for 1 hour. Nematodes were imaged at a confocal Zeiss LSM880 microscope with a 20x objective at indicated time points after HS. Animals were scored for clearly visible nuclear or cytosolic localization of GFP. Statistical analysis was performed using a Kruskal-Wallis test.

### Western Blot analysis

Protein extracts were prepared from age-synchronized 1-day adult nematodes grown in liquid culture or on 90 mm NGM plates at a population density of 1000 nematodes per plate. Nematodes were harvested into a pellet by washing three times in M9 buffer and the pellet was flash-frozen in liquid nitrogen. The frozen worm pellet was ground with an Eppendorf pestle and resuspended in Lysis Buffer (10 mM Tris pH 7.5; 150 mM NaCl; 0.5 mM EDTA; 0.5% NP-40), supplemented with protease inhibitor cocktail (Complete Mini, EDTA-free, Roche) and phosphatase inhibitor (PhosSTOP, Roche). The cell extract was then prepared by centrifugation at 10,000 x g for 5 minutes at 4°C and protein concentration was determined by Bio Rad protein assay kit (Bradford assay). Cell extracts were mixed with 5x SDS sample buffer and boiled for 5 minutes. 25 ug total protein as loaded onto a 10 % SDS PAGE and Western blot analysis was performed as described previously (van Oosten-Hawle et al., 2013). An anti-Flag antibody (Sigma) was used to detect FLAG-tagged fusion proteins (PQM-1 and DAF-16), and anti-tubulin antibody (Sigma, mouse monoclonal) was used to detect tubulin as a loading control.

### Quantitative Real-time PCR

RNA was extracted using TriZOL reagent. A pellet grinder was used to grind the frozen *C. elegans* pellet on ice 3x for 30 seconds. RNA was purified using the Zymo-prep RNA Mini Isolation kit (Zymo Research, Cambridge Biosciences) and qRT-PCR was performed as described previously (O’Brien et al., 2018; van Oosten-Hawle et al., 2013). RNA concentration was measured using a Thermo Scientific NanoDrop One, and 100 ng of RNA was reverse transcribed into cDNA using a Bio-Rad iScript cDNA synthesis kit. Quantitative PCR was performed using Bio-Rad Universal SYBR green Supermix in a Bio-Rad CFX Connect Real-Time System. 3 biological replicates were performed per sample. Significance was determined using Student’s t test or one-way ANOVA with a cutoff of p < 0.05.

### Heat stress survival assays

Synchronized populations of approximately 100 L4-staged nematodes were grown on a 35 mm NGM plate seeded with OP50-1 *E. coli.* Plates were incubated in a water bath equilibrated at 35°C for 4 hrs. Nematodes were allowed to recover for at least 16 hours at 20°C before scoring for touch induced movement and pharyngeal pumping. Each experiment was repeated at least three times.

### Oxidative stress survival assays

Synchronized populations of approximately 200 L4-staged nematodes were grown on a 60 mm NGM plate, seeded with OP50-1 *E. coli*. Nematodes were washed off the plates in M9 buffer and incubated in 150 mM paraquat, containing *E. coli*, at a density of one nematode per uL for 6 hrs. Nematodes were gently washed three times in M9 buffer and allowed to recover on 35 mm plates seeded with OP50-1 *E. coli*. Sixteen hours later they were scored for touch induced movement and pharyngeal pumping. Each experiment was repeated at least five times.

### Dietary restriction

Nematodes were grown until Day-1 adulthood, then collected and washed three times in M9 buffer, and plated on NGM Agar plates devoid of bacterial food. After indicated time periods (2, 4, 6 and 24 hrs) animals were collected by washing off with M9 buffer and flash freezing in liquid nitrogen before further processing for qRT-PCR or Western blot analysis

### Statistical Analysis

All experiments were repeated at least three times (3 biological replicates). Data are presented as mean values ± SEM. p values were calculated using Student’s t test. ∗ denotes p < 0.05; ∗∗ denotes p < 0.01, and ∗∗∗ denotes p < 0.001; (ns) denotes non-significant. Statistical analysis was carried out using Graph Pad Prism Software (version 9).

### Data and Software availability

The data discussed in this study have been deposited in NCBI’s Gene Expression Omnibus and are accessible through GEO Series accession number **GSE287678.**

### Transcriptomic profiling via RNA-seq

RNA was extracted from samples as described above. Agarose gel electrophoresis using a 1% gel was performed for a visual determination of sample quality, and RNA integrity number (RIN) was determined by the University of Leeds Next Generation Sequencing Facility using an Agilent 2200 TapeStation. RNA-seq was performed by Novogene on an Illumina Hi-Seq platform. Calculation of log_2_(fold change), p values and corrected p values were performed by Novogene. WBCel235 was used as the reference genome for annotation. GO term and KEGG pathway analysis was performed using the publicly available online tool DAVID Bioinformatics Resources 6.8.

## Supporting information

Supplemental files combined

## Supplemental Figure legends

**Supplemental Figure 1. PQM-1 expression and nuclear localization dynamics during heat- and oxidative stress.**

**(A)** Confocal images of PQM-1::GFP expression in L4 larvae (strain OP201) before HS (−1 hr), immediately after a 1-hour HS at 35°C (0 hr) and after 2 hours of recovery post HS (+ 2hr). PQM-1::GFP is localized to intestinal nuclei in L4 stage larvae during normal growth conditions.

**(B)** Confocal images of Day-1 adult PQM-1::GFP::FLAG expression after (0 hr, 2 hr and 4 hr) oxidative stress (150 mM paraquat). **(C)** Western blot analysis of PQM-1::GFP::FLAG expression immediately after (0 hr) and 1 – 4 hrs post oxidative stress, using an anti-FLAG antibody and tubulin as a loading control. **(D)** *pqm-1* transcript levels before (−1 hr) and 0, 1, 2, 4, 6 hours after a 1-hr exposure to oxidative stress (150 mM paraquat).

**(E)** Survival to oxidative stress of Day 1 adults grown on control or *skn-1* RNAi and treated with 150 mM Paraquat for 6 hours. n = 20; three biological replicates. Bar graph represent SEM, *P < 0.05; n.s. = not significant.

**Supplemental Figure 2. PQM-1 nuclear localization and transcript expression before and after HS.**

**(A)** Confocal images of PQM-1::GFP expression in Day 1 adult nematodes before HS (− 1hr), immediately after a 1-hour 35C HS (0 hr) and 1, 2, 4 and 6-hours post HS. Scale bar = 50 μm.

**(B)** Nuclear PQM-1::GFP fluorescence intensity before and after HS in Day-1 adults. **(C)** *pqm-1* transcript levels before (−1 hr) and 0, 1, 2, 4, 6 hours after HS.

**Supplemental Figure 3. RNA-Seq analysis of *pqm-1* dependent gene expression in heat-shocked Day 1 adults.**

**(A)** RNA-Seq scatterplot showing log2 fold-changed expression levels of differentially expressed genes (P value < 0.05) 2 hours after HS in control (N2) worms **(B)** Scatterplot showing log2 fold-changed expression levels of differentially expressed genes (P-Value < 0.05) in *pqm-1(ko)* mutants 2 hours post HS. **(C)** Scatterplot of log2 fold-changed gene expression levels (P < 0.05) in *daf-16* mutants 2 hours post HS.

**(D)** Venn diagram analysis of chaperone genes upregulated 2 hours post HS (P_adj_ < 0.1) in wild type (“*pqm-1(+);daf-16(+)*”); *pqm-1(k.o)* mutant (“*pqm-1(−);daf-16(+))* and *daf-16(mu86)* mutant (*pqm-1(+);daf-16(−)*. **(E)** 33 chaperone genes upregulated 2 hours post-HS in wild type animals (*pqm-1(+);daf-16(+)*), exclusively dependent on *pqm-1*, and exclusively dependent on *daf-16*.|

**(F)** RNAi mini screen to identify requirement of the 42 PQM-1-dependent genes for heat stress survival. **(G)** Transcript expression of identified 13 *pqm-1* dependent candidate genes required for heat stress survival in control (N2), *pqm-1(ko)* mutant and *pqm-1 (phospho-null)* mutant before and after HS (1h, 35°C). n=3. Error bars represent S.E.M. One-way ANOVA. *P<0.05; **P<0.01; ***P<0.001; ****P<0.0001; n.s. = not significant.

## Acknowledgments

This work was supported by start-up funds by the University of North Carolina at Charlotte and an NIH grant R01AG082970 to P.v.O.H. We acknowledge support by the NC3Rs (NC/P001203/1) to L.M.J., awarded to P.v.O.H, and an NIGMS grant R35GM137985 to R.H.D. Some *C. elegans* strains used in this research were provided by the Caenorhabditis Genetics Center, which is funded by the NIH Office of Research Infrastructure Programs (P40 OD010440).

