## Supplemental files combined for "Stress-dependent activation of PQM-1 orchestrates a second-wave proteostasis response for organismal survival"

### Supplemental Figure 1

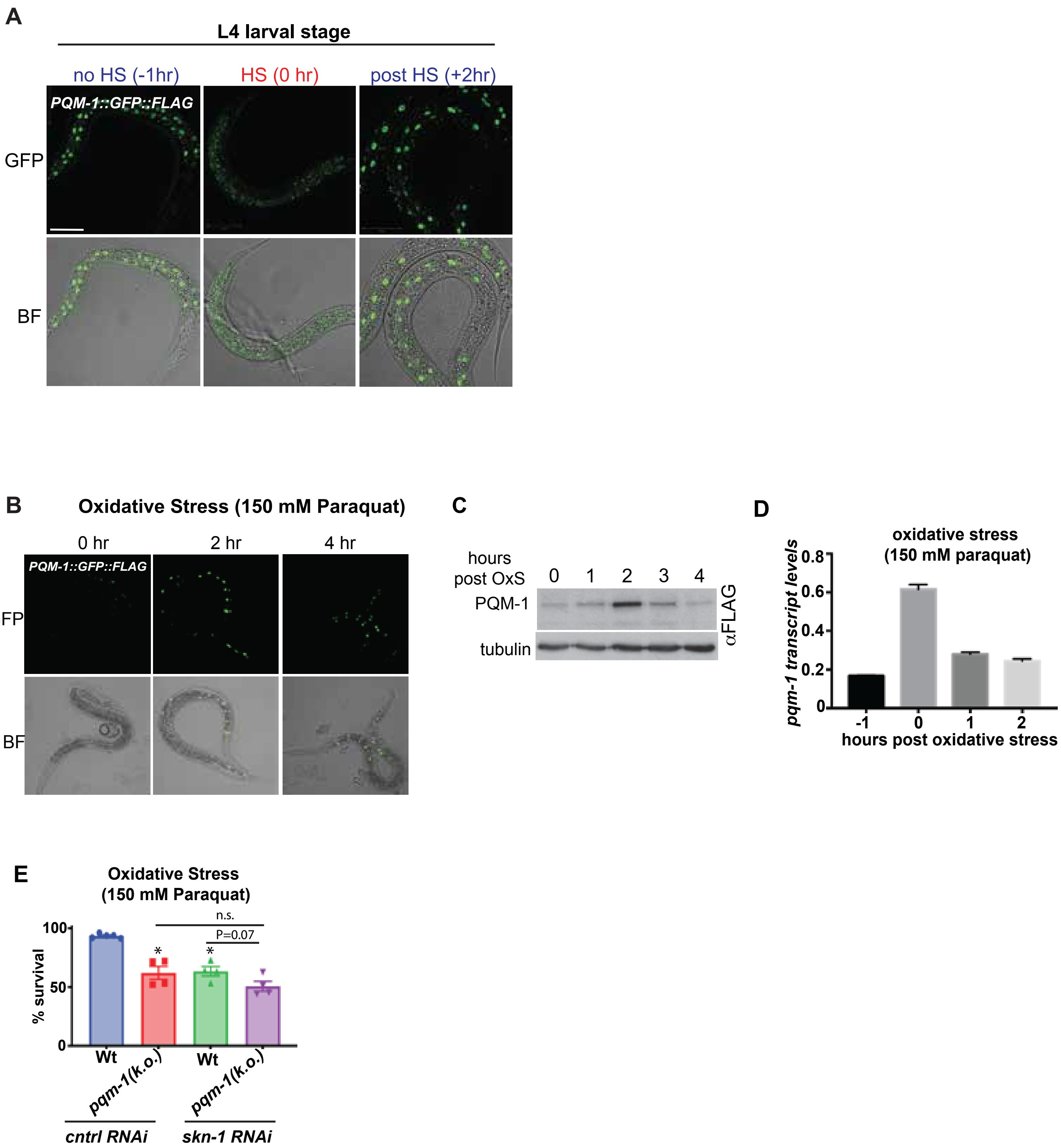

Supplemental Figure 2

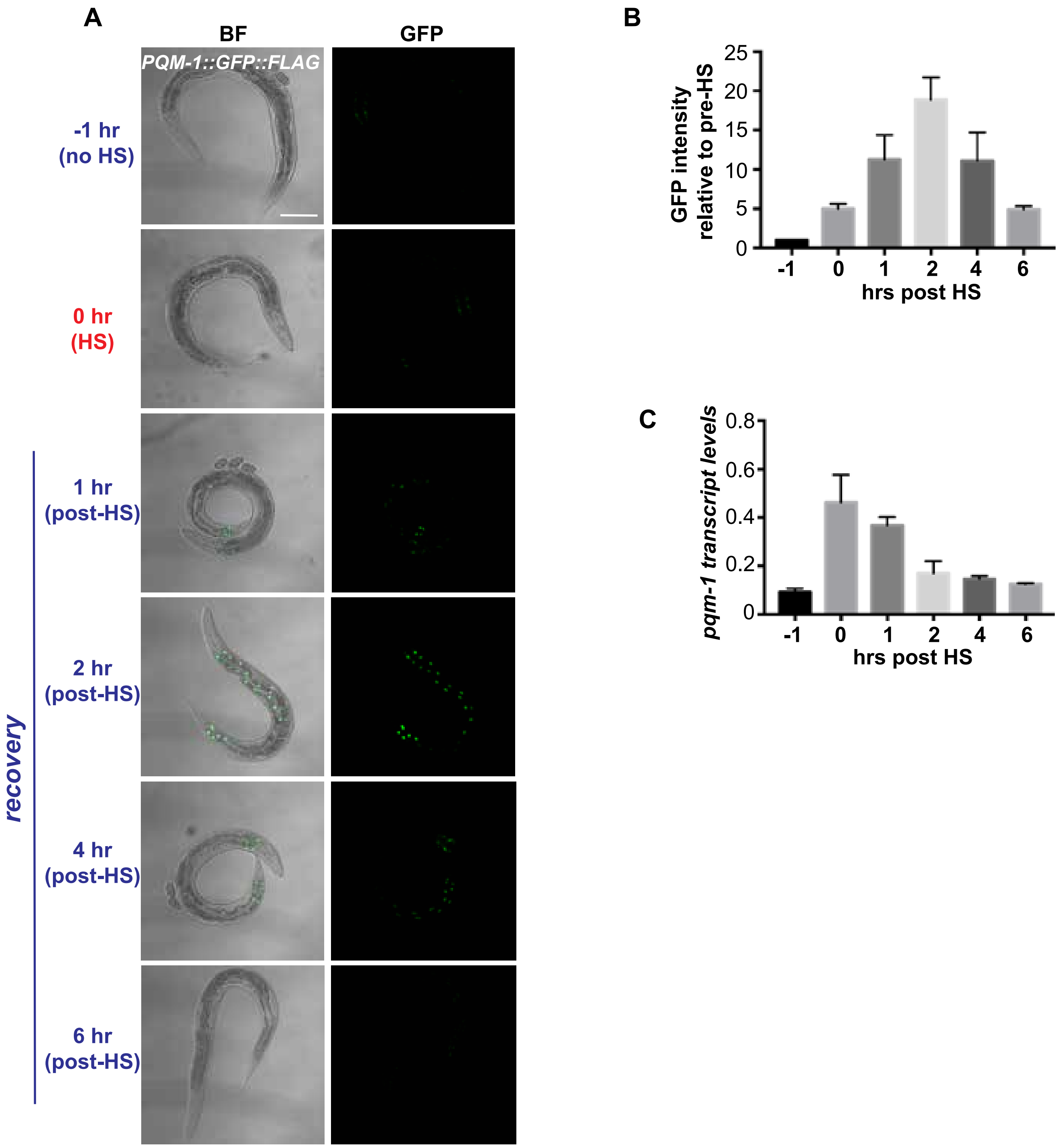

Supp Figure 3

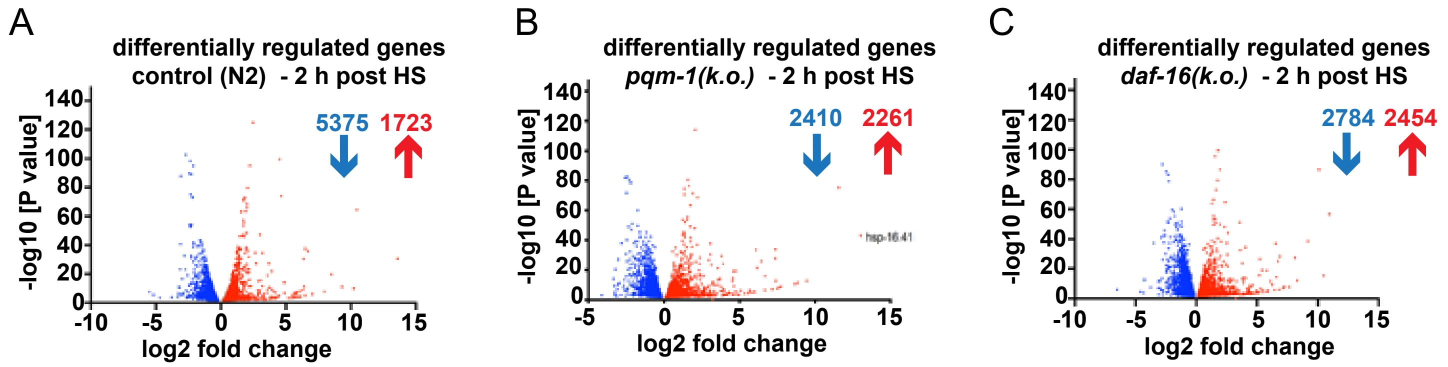


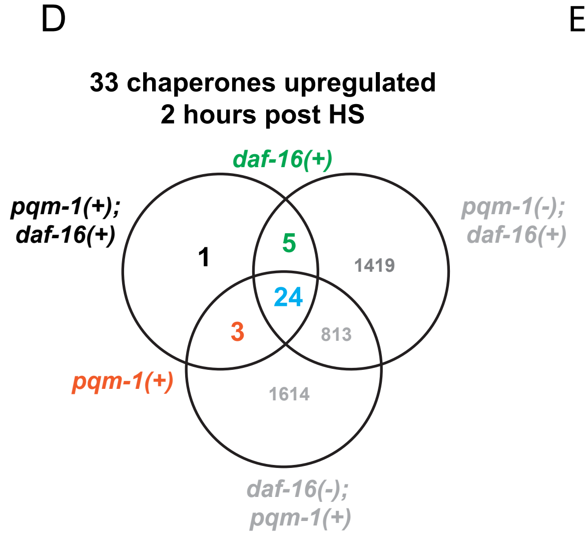


E

| Chaperone Family | C. elegans chaperone | daf-16(+) | pqm-1(+) |
|---|---|---|---|
| Hsp70 | T14G8.3 | T14G8.3 | |
| Hsp70 | F44E5.4 | F44E5.4 | |
| Hsp70 | F44E5.5 | F44E5.5 | |
| Hsp70 | hsp-4 | | |
| Hsp70 | hsp-70 | | |
| Hsp70 | hsp-1 | | hsp-1 |
| Hsp90 | hsp-90 | | |
| Hsp40 | dnj-7 | | dnj-7 |
| Hsp40 | dnj-13 | | |
| Hsp40 | dnj-27 | | |
| Hsp40 | dnj-14 | | |
| Hsp40 | dnj-12 | | |
| Hsp70/Hsp90 cochap | F52H5.3 | | |
| Hsp70/Hsp90 cochap | dpy-18 | | dpy-18 |
| Hsp70/Hsp90 cochap | ogt-1 | | |
| Hsp70/Hsp90 cochap | klc-2 | | |
| Hsp70/Hsp90 cochap | sgt-1 | | |
| Hsp70/Hsp90 cochap | prx-5 | | |
| Hsp70/Hsp90 cochap | Y73F8A.26 | Y73F8A.26 | |
| sHSP | F08H9.4 | F08H9.4 | |
| sHSP | hsp-16.2 | | |
| sHSP | hsp-16.41 | | |
| sHSP | hsp-43 | | |
| sHSP | hsp-12.6 | | |
| sHSP | hsp-25 | | |
| Hsp90 cochap | unc-45 | | |
| Hsp90 cochap | cdc-37 | | |
| Hsp90 cochap | cyn-8 | | |
| Hsp90 cochap | fkb-6 | | |
| Hsp90 cochap | fkb-4 | | |
| Hsp60/Chaperonin | cct-7 | | |
| Hsp60/Chaperonin | cct-3 | | |
| Hsp60/Chaperonin | cct-5 | | |

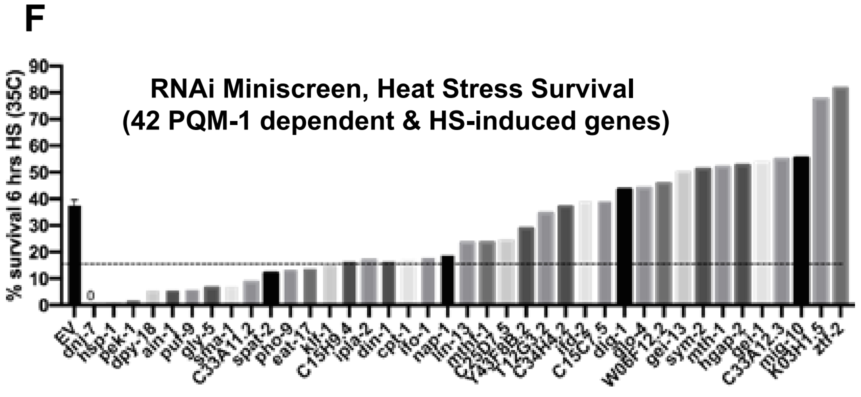


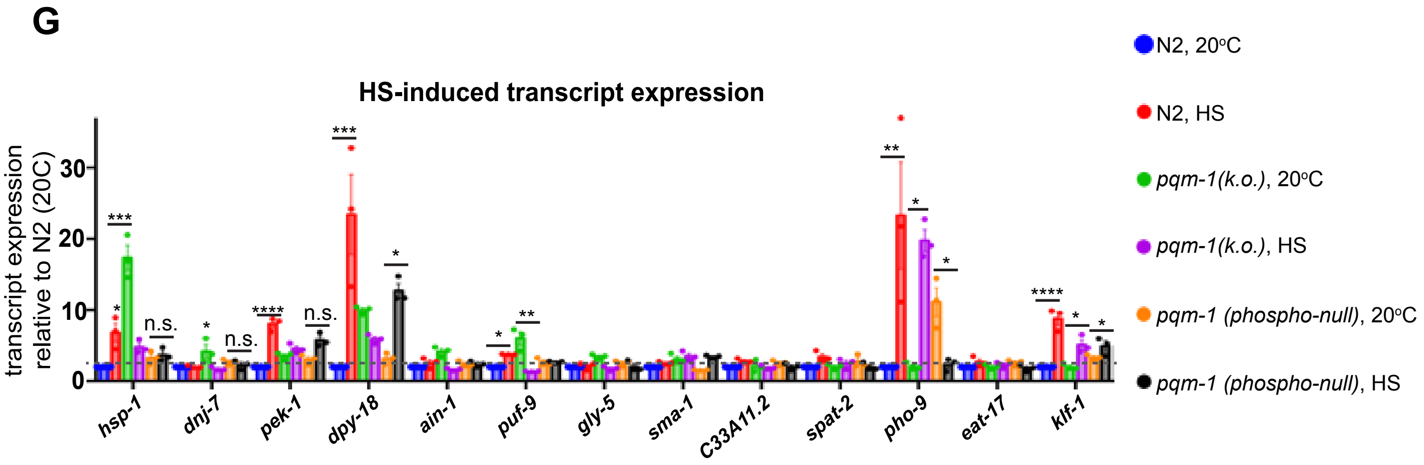
